# A highly expressing, soluble, and stable plant-made IgG fusion vaccine strategy enhances antigen immunogenicity in mice without adjuvant

**DOI:** 10.1101/2020.07.13.199703

**Authors:** Andrew G. Diamos, Mary D. Pardhe, Haiyan Sun, Joseph G. L. Hunter, Jacquelyn Kilbourne, Qiang Chen, Hugh S. Mason

## Abstract

Therapeutics based on fusing a protein of interest to the IgG Fc domain have been enormously successful, though fewer studies have investigated the vaccine potential of IgG fusions. In this study, we systematically compared the key properties of seven different plant-made human IgG1 fusion vaccine candidates using Zika virus (ZIKV) envelope domain III (ZE3) as a model antigen. Complement protein C1q binding of the IgG fusions was enhanced by: 1) antigen fusion to the IgG N-terminus; 2) removal of the IgG light chain or Fab regions; 3) addition of hexamer-inducing mutations in the IgG Fc; 4) adding a self-binding epitope tag to create recombinant immune complexes (RIC); or 5) producing IgG fusions in plants that lack plant-specific β1,2-linked xylose and α1,3-linked fucose N-linked glycans. We also characterized the expression, solubility, and stability of the IgG fusions. By optimizing immune complex formation, a potently immunogenic vaccine candidate with improved solubility and high stability was produced at 1.5 mg IgG fusion per g leaf fresh weight. In mice, the IgG fusions elicited high titers of Zika-specific antibodies which neutralized ZIKV using only two doses without adjuvant, reaching up to 150-fold higher antibody titers than ZE3 antigen alone. We anticipate these findings will be broadly applicable to the creation of other vaccines and antibody-based therapeutics.

**Highlights:**

- A modified immune complex has high expression, solubility, stability, and immunogenicity.
- Antigen immunogenicity is improved up to 150-fold by fusion to plant-made IgGs.
- High serum IgG titers >1:500,000 were achieved with only two doses without adjuvant.

## 1. Introduction

Subunit vaccines consisting of recombinant protein antigens are very promising due to their safety, ease of production, and capacity to elicit targeted immune responses directed towards desired epitopes. However, when delivered by themselves, these antigens often fail to generate robust and long-lasting immune responses, necessitating strategies to enhance their immunogenicity (Reed et al., 2013). Protein fusions to a variety of bioengineered IgG-based scaffolds have demonstrated tremendous potential as therapeutic candidates, acting to enhance the solubility and stability of the fusion partner while also allowing simple and cost-effective purification via protein A/G affinity chromatography (Carter, 2011). Furthermore, by interacting with neonatal Fc receptors (FcRn) in the body, IgG fusions can escape lysosomal degradation, thereby extending the serum half-life of the Fc-fusion (Roopenian and Akilesh, 2007).

Though less well explored, IgG fusions also have many desirable characteristics for vaccine development. Antigen-presenting cells can uptake and process IgG-bound antigen via FcRn receptors (Baker et al., 2011; Qiao et al., 2008), complement receptor C1q (Fletcher et al., 2018; McCloskey et al., 2011; West et al., 2018) and Fcγ receptors (Bournazos and Ravetch, 2017). However, not all IgG subclasses interact equally with immune receptors. IgG1 and IgG3 are immunostimulatory, while IgG2 and IgG4 have weak effector functions (Vidarsson et al., 2014). IgG3 mediates the strongest complement activation and Fc receptor binding, however it has significantly reduced serum half-life, and thus most IgG-fusions vaccine strategies use human IgG1 (Czajkowsky et al., 2012). In addition to these factors, the polymeric state of an antibody also affects its immunological properties. Both C1q, the first component involved in complement activation, and the low affinity Fcγ receptors require high avidity binding for activation. For this reason, monovalent Fc-antigen fusions cannot efficiently utilize these pathways. By contrast, larger antigen-antibody immune complexes with multivalent Fc domains can efficiently bind C1q and cross-link Fc receptors, resulting in greatly improved uptake and presentation by dendritic cells, and subsequently improved activation of T-cells (Fletcher et al., 2018; Getahun and Heyman, 2006; Ho et al., 2017; van Montfoort et al., 2012).

Capitalizing on the benefits of antigen-antibody immune complexes, a variety of vaccine strategies have been developed to improve antigen immunogenicity. Immune complexes have been used to focus the immune response towards favorable antigenic sites (Hioe et al., 2018; Tsouchnikas et al., 2015) as well as enhance the potency and breadth of the immune response by selecting for high affinity B-cells (Maamary et al., 2017). By engineering antigen-IgG fusions to form polymeric structures via inclusion of the IgM tailpiece, both humoral and cellular immune responses were enhanced in the absence of adjuvant (Kim et al., 2018; Webster et al., 2018). While traditional immune complexes consisting of antigen mixed with antibody will form variable structures depending on the oligomeric status of the antigen, we and others have developed a recombinant immune complex (RIC) strategy that allows consistent formation of large immune complexes which can efficiently engage complement (Mason, 2016). In the RIC system, a well-characterized antibody is tagged with its own binding site and is fused to an antigen of choice. The resulting molecules self-interact and form highly immunogenic antigen-antibody clusters. Past research has shown the potential for RIC-like molecules to produce promising vaccine candidates for *Clostridium tetani* (Chargelegue et al., 2005), Ebola virus (Phoolcharoen et al., 2011a, 2011b), *Mycobacterium tuberculosis* (Pepponi et al., 2014), dengue virus (Kim et al., 2015), human papillomavirus (Diamos et al., 2019) and Zika virus (Diamos et al., 2020b).

Zika virus (ZIKV) envelope domain III (ZE3) is a promising subunit vaccine candidate due to its ability to generate neutralizing antibodies without potential for antibody-dependent enhancement (Shukla et al., 2020; Stettler et al., 2016; Yang et al., 2017). However, like many subunit vaccines consisting of viral antigens, it is not strongly immunogenic on its own, necessitating high doses with adjuvant and repeated immunizations (Yang et al., 2017). We have previously shown that RIC carrying ZE3 elicits strong immune responses (Diamos et al., 2020b). However, traditional RIC may be undesirable, as the large complex size renders them poorly soluble and limits their viability as a vaccine candidate (Diamos *et al.*, 2019). Therefore, in this study we built on recent advances in antibody engineering to design and explore a panel of modified IgG fusion vaccines carrying ZE3 as a model antigen. These constructs contained modifications designed to alter their polymeric and C1q binding characteristics. We then compared the C1q binding, expression, solubility, stability, and immunogenicity of the IgG fusions.

## 2. Results

### 2.1 Modified plant-made IgG-ZE3 fusions exhibit variable binding to complement receptor C1q

We designed a panel of human IgG1 variants based on the previously characterized humanized anti-Ebola monoclonal antibody 6D8 (Phoolcharoen et al., 2011a) fused to ZE3 (**Fig. 1A**). A RIC construct was created by fusing 6D8 epitope-tagged ZE3 via a flexible linker to the C-terminus of the 6D8 heavy chain. When in solution, the 6D8 Fab arms of one 6D8-ZE3 fusion molecule bind the epitope-tagged ZE3 on another fusion molecule, forming antigen-antibody clusters (Mason, 2016). This RIC construct is referred to as "HLZe" as it contains the 6D8 **H**eavy chain, 6D8 **L**ight chain, C-terminal **Z**E3 fusion, and 6D8 **e**pitope binding tag. As a control for immune complex formation, an otherwise identical construct was created without the epitope tag (construct “HLZ”). This construct would not be expected to form immune complexes as it cannot specifically bind other HLZ molecules. The 6D8 antibody without ZE3 or epitope tag is referred to as construct “HL,” and the corresponding RIC with epitope tag but lacking ZE3 is referred to as “HLe.” Since C1q mediates immune complex uptake into antigen presenting cells and complement-coated immune complexes play a key role in activating and maintaining long-term immunity (Croix et al., 1996; McCloskey et al., 2011; Phan et al., 2007; West et al., 2018), the constructs were analyzed for C1q binding. Additionally, C1q binding provides insight into the polymeric status of the constructs, as RIC and other polymeric IgGs have stronger binding to C1q when compared with monomeric IgG (Chargelegue et al., 2005; Webster et al., 2018). To improve C1q binding, all constructs were expressed in glycoengineered *Nicotiana benthamiana* that have suppressed fucosyl-and xylosyl transferase activities, as these glycans inhibit immune receptor binding (Kallolimath and Steinkellner, 2015). HL or HLe made in glycoengineered plants showed highly improved C1q binding compared to constructs expressed in wildtype plants (**Fig. S1A**). Consistent with the formation of densely clustered antigen-antibody complexes, HLZe showed greatly improved C1q binding compared to HLZ (p < 0.001) (**Fig. 1B**). A small increase in C1q binding was noted with HLZ compared to HL (p < 0.05) (**Fig. 1B**), suggesting low-level aggregation of the construct or slight favorable alteration of Fc conformation due to the ZE3 fusion. HLZe also efficiently bound FcγRIIIa, exceeding the binding of HLZ and HL (**Fig. S1B**).

**Figure 1.**
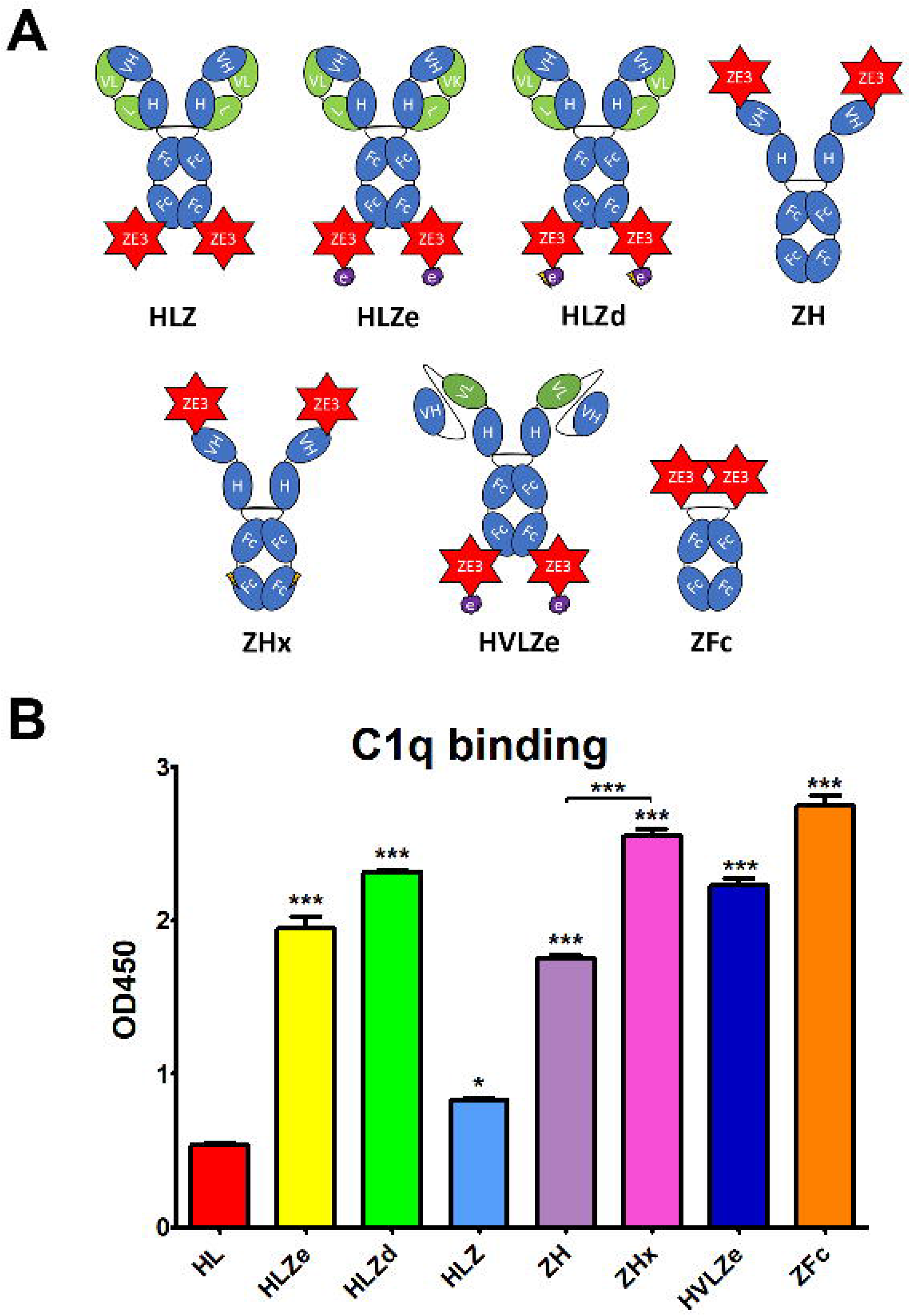
Various IgG fusion strategies enhance C1q binding. (A) Schematic representation of IgG fusion constructs used in this study. Fusion constructs are built on the mAb 6D8 human IgG1 backbone with the shown modifications. ZE3; the Zika envelope domain III containing amino acids K301 to T406; e, an epitope tag containing the “VYKLDISEA” 6D8 binding motif for RIC formation; e with lightning bolt, an epitope tag truncated to include only “YKLDIS” to reduce RIC formation; VH, the variable heavy domain from 6D8 which participates in binding the epitope tag; VL, the variably light domain from 6D8 which participates in binding the epitope tag; H, the heavy chain constant CH1 domain from 6D8; L, the light chain constant domain from 6D8; Fc, the heavy chain constant CH2 or CH3 domains from 6D8; Fc with lightning bolt, same as Fc but with E345R, E430G, and S440Y mutations to induce hexamer formation. (B) C1q binding ELISA of purified IgG fusion constructs made in plants that lack plant-specific β1,2-linked xylose and α1,3-linked fucose N-linked glycans. ELISA plates were coated with 10 μg/ml human C1q and incubated with 10 μg/ml each molecule, using 6D8 with no fusion as a negative control. Constructs were detected using polyclonal goat anti-human IgG-HRP. Mean OD_450_ values from three samples are shown ± standard error with one star (*) indicating p < 0.05 and three stars (***) indicating p < 0.001 as measured by one-way ANOVA comparing each column to the control column HL, unless otherwise indicated with a bracket.

Hexamers are the ideal substrate for C1q binding (Diebolder et al., 2014) and it has been suggested that the antibody Fab arms play a regulatory role in complement activation by inhibiting C1q binding (Wang et al., 2016). Upon addition of certain soluble antigens or removal of the light chains, C1q binding is dramatically improved, suggesting conformational changes are conveyed into the Fc domain from the Fab arms to induce hexamer formation (Wang et al., 2016). In agreement with this model, we find that the addition of soluble antigen carrying the 6D8 epitope tag modestly improved C1q binding (**Fig. S2**, compare HL vs HL + Ag) and C1q binding was strongly enhanced by removal of the antibody light chain (**Fig. S2**, compare HL and H). Interestingly, antigen fusion to the heavy chain N-terminus also strongly improved C1q binding at a level similar to removing the light chain (**Fig. S2**, compare HL, H, and ZHL). Therefore, we created a construct with ZE3 fused to the N-terminus of the heavy chain (construct “ZH”) in the absence of the light chain. ZH efficiently bound C1q similar to HLZe; however, the effects of light chain removal and N-terminal ZE3 fusions were not additive (**Fig. 1B**). In an attempt to further improve interaction with C1q, we made a construct that was similar to ZH except for mutations E345R, E430G, and S440Y in the Fc region which favor formation of hexamers (Diebolder et al., 2014). This construct (ZHx) had modestly improved C1q binding compared to ZH (p < 0.001) (**Fig. 1B**). Past studies have shown that IgG1 Fc fusions (lacking VH and CH1 domains) produce variable levels of C1q binding depending on the fusion partner (Lagassé et al., 2019) . However, Fab arm removal can favor hexamer formation with strong complement activation (Wang et al., 2016). ZE3 N-terminally fused to the 6D8 Fc (**Fig. 1A**construct “ZFc”) showed very strong C1q binding (**Fig. 1B**). Finally, as RIC require co-expression of heavy and light chains, we constructed a simplified RIC that consisted of a single chain antibody with the variable light (VL) domain of 6D8 inserted between the variable heavy (VH) and CH1 domains of 6D8 to yield construct “HVL”. This construct bound antigen tagged with the 6D8 epitope, albeit at a somewhat reduced level compared to unaltered 6D8 (**Fig. S3**). This single-chain Fab configuration was fused to ZE3 to create the single chain RIC HVLZe (**Fig. 1A**), which displayed strong C1q binding (**Fig. 1B**). Together, these data demonstrate that RIC formation, removal or rearrangement of light chain/Fab region, Fc hexamer mutations, and ZE3 fusion to the IgG N-terminus all increase C1q binding of plant-made IgG1.

### 2.2 Reducing RIC self-binding improves solubility without reducing C1q binding

RIC suffer from low yield of soluble product (Diamos et al., 2019). While unmodified 6D8 antibody is highly soluble, addition of the short epitope tag (construct HLe) renders the antibody mostly insoluble (**Fig. S4**). However, this is prevented by removal of the light chain (construct He), which is needed for epitope binding (Kim et al., 2015), suggesting that the insolubility arises from large complexes of antibody bound to the epitope tag (**Fig. S4**). To improve RIC solubility, we shortened the 6D8 epitope tag in order to reduce the strength of self-binding. Construct “HLa” contains the minimal reported binding region for 6D8 “VYKLDISEA” (Wilson et al., 2000). Removal of a single amino acid from the C-terminus (construct “HLb,” epitope VYKLDISE) had no effect on solubility (**Fig. S5A**). However, further removal of a single amino acid from the N-terminus (construct “HLc,” YKLDISE), and additional truncation in construct “HLd” (YKLDIS) resulted in greatly improved solubility, yielding 20-fold more soluble antibody (**Fig. S5B**). Thus, the truncated epitope present in HLd was introduced to HLZe (construct HLZd) and characterized. Despite reducing epitope binding compared to full length epitope tag by approximately 25-fold by ELISA (**Fig. S5C**), HLZd still maintained very strong C1q binding, suggesting efficient complex formation still remained (**Fig. 1B**). Note that the otherwise identical construct HLZ, which lacks the YKLDIS tag, had 10-fold lower C1q binding (**Fig. S5D**). These data suggest that RIC solubility can be enhanced by reducing self-binding without a loss of C1q activation.

### 2.3 Highly variable expression and solubility of IgG fusions in plants

Since a viable vaccine candidate must be soluble and highly expressing, we measured the soluble yield of fully assembled IgG fusions by employing an ELISA assay that first captured ZE3 and then detected human IgG. As a standard, purified and quantified HLZ was used. To detect any proteolytic cleavage of ZE3, an ELISA measuring total IgG irrespective of the presence of ZE3 was also used as a comparison, using purified and quantified unfused HL as standard. When soluble extracts were probed for the presence of both ZE3 and IgG, construct ZH yielded 0.83 mg fully formed product per gram leaf fresh weight (mg/g LFW), and the similar ZFc yielded 0.58 mg/g LFW (**Fig. 2A**). However, when measuring the total IgG content, the yield of ZH was roughly 1.2-fold higher and, for ZFc, 2.5-fold higher than ZE3 ELISA, which suggests that ZH and especially ZFc are susceptible to proteolytic cleavage. This finding was confirmed by visualization of the ZFc cleavage products on SDS-PAGE (**Fig. 2B**). HLZ accumulated only 0.17 mg/g LFW fully assembled product, with roughly 40% ZE3 lost to cleavage, suggesting general instability of the construct (**Fig. 2A, 2B**). By ELISA, the polymeric RIC constructs seemingly had lower yields than the monomeric constructs: HLZe and HLZd only accumulated 0.04 mg/g LFW and 0.30 mg/g LFW respectively, and the hexameric ZHx accumulated only 0.08 mg/g LFW (**Fig. 2A**). However, when visualized under reducing SDS-PAGE conditions, HLZd was the highest expressing construct, accumulating an estimated 1.5 mg/g LFW, over twice the highest yield of the next best construct, with no visible degradation products (**Fig. 2A, 2B**). The discrepancy between ELISA and gel quantification likely arises due to complexed HLZd being less accessible to the antibody probe by ELISA. Importantly, the gel quantification results agreed with the ELISA results for all the monomeric constructs, but not for any of the polymeric constructs (**Fig. 2A**). Thus, the total yield of HLZe and ZHx was also higher when measured by gel quantification (**Fig. 2A, 2B**). While polymeric constructs were partially inaccessible to the antibody probe by traditional ELISA (**Fig. 2A**), they were strongly detected using the same antibody probe in the C1q assay (**Fig. 1B**). This likely occurred due to C1q disaggregating the polymeric constructs, as C1q functions to solubilize immune complexes *in vivo* (Thielens et al., 2017). Construct HVLZe yielded only 0.02 mg/g LFW soluble product by ELISA and was not visible by SDS-PAGE (**Fig. 2A**). However, we detected very high levels when extracting with 7.5 M urea (data not shown) suggesting that the construct insolubility resulted in the low yield. In total, these results show that the different IgG fusions display expression levels which vary by several orders of magnitude, highlighting the large differences in their capacity to be used as viable, affordable vaccines.

**Figure 2.**
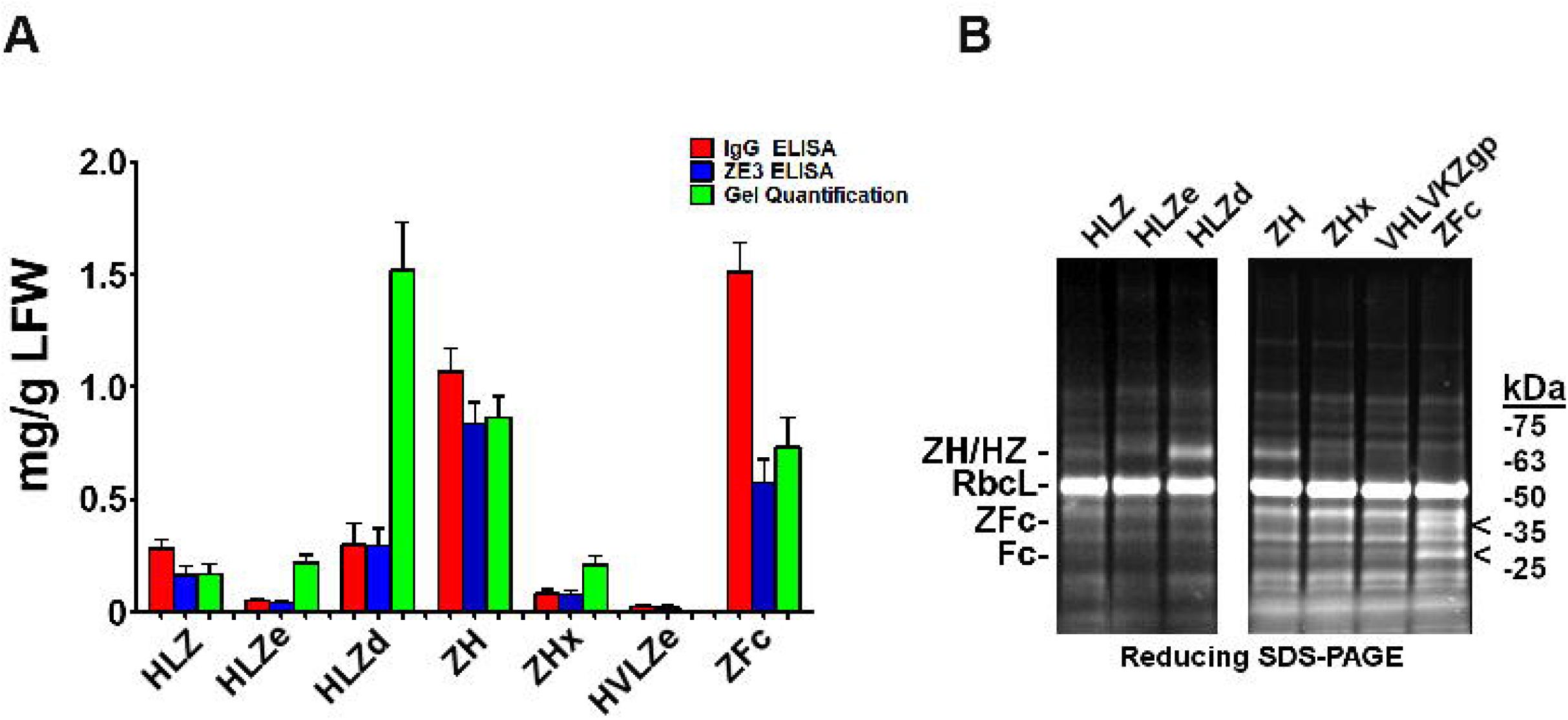
Modified IgG fusions have improved expression and solubility. (A) ELISA and gel quantification of IgG fusion construct expression. Clarified protein extracts (“soluble” fraction) from leaf spots agroinfiltrated with each IgG fusion construct were analyzed by either ELISA, or SDS-PAGE followed by gel image quantification. For ZE3 ELISA, plates were coated with polyclonal mouse anti-ZE3, incubated with serial dilutions of extracts from each IgG fusion using purified HLZ as a standard, and probed with goat anti-human IgG-HRP. For IgG ELISA, plates were coated with serial dilutions of extracts or human IgG standard and probed with goat anti-human IgG-HRP. For gel quantification, ImageJ software was used to compare the IgG fusion band intensity visualized on stain-free polyacrylamide gels using purified 6D8 antibody as standard. Columns represent means ± standard error from three independently infiltrated leaf samples. (B) Clarified leaf extracts were separated by reducing SDS-PAGE and a representative gel image is shown. The general band position corresponding to each respective heavy chain/ZE3 fusion is indicated “ZH/HZ.” The small shift in size in HLZe (~67 kDa of the reduced heavy chain fusion before glycosylation) and HLZd (~65 kDa) or HLZ (~64.5 kDa) is due to epitope tag truncation. ZH and ZHx have a predicted size of ~64.5 kDa. The “<” indicates ZE3-Fc (~40 kDa) and Fc (~29 kDa) fragments. The large subunit of Rubisco is abbreviated “RbcL.”

### 2.4 Purification and aggregation of IgG fusion constructs

IgG fusions were purified to >95% homogeneity using a simple one-step purification via protein G affinity chromatography. In agreement with the expression data, the more polymeric constructs showed less degradation than the other constructs, and the ZFc fusion had particularly high levels of degradation (**Fig. 3**). To investigate the polymeric characteristics of each construct, purified IgG fusions were analyzed by sucrose gradient sedimentation. Consistent with the formation of large immune complexes, HLZe and HVLZe were found mostly in the bottom of the gradient while HLZ, ZH, and ZFc were found mostly at the top of the gradient (**Fig. 4A, 4B**). A small shift in density was noted with construct HLZ (**Fig. 4A**) compared to unfused HL, which may explain the increased C1q binding (**Fig. 1B**). Soluble extracts of ZHx contained both low-density material as well as some very high-density material, suggesting that the combination of light chain removal and RGY mutations contributes to the formation of larger aggregates (**Fig. 4B**). Extracts of HLZd showed intermediate density compared to HLZ and HLZe (**Fig. 4A**) which, when taken together with the expression data and C1q binding, are consistent with HLZd forming smaller, more soluble immune complexes compared to HLZe.

**Figure 3.**
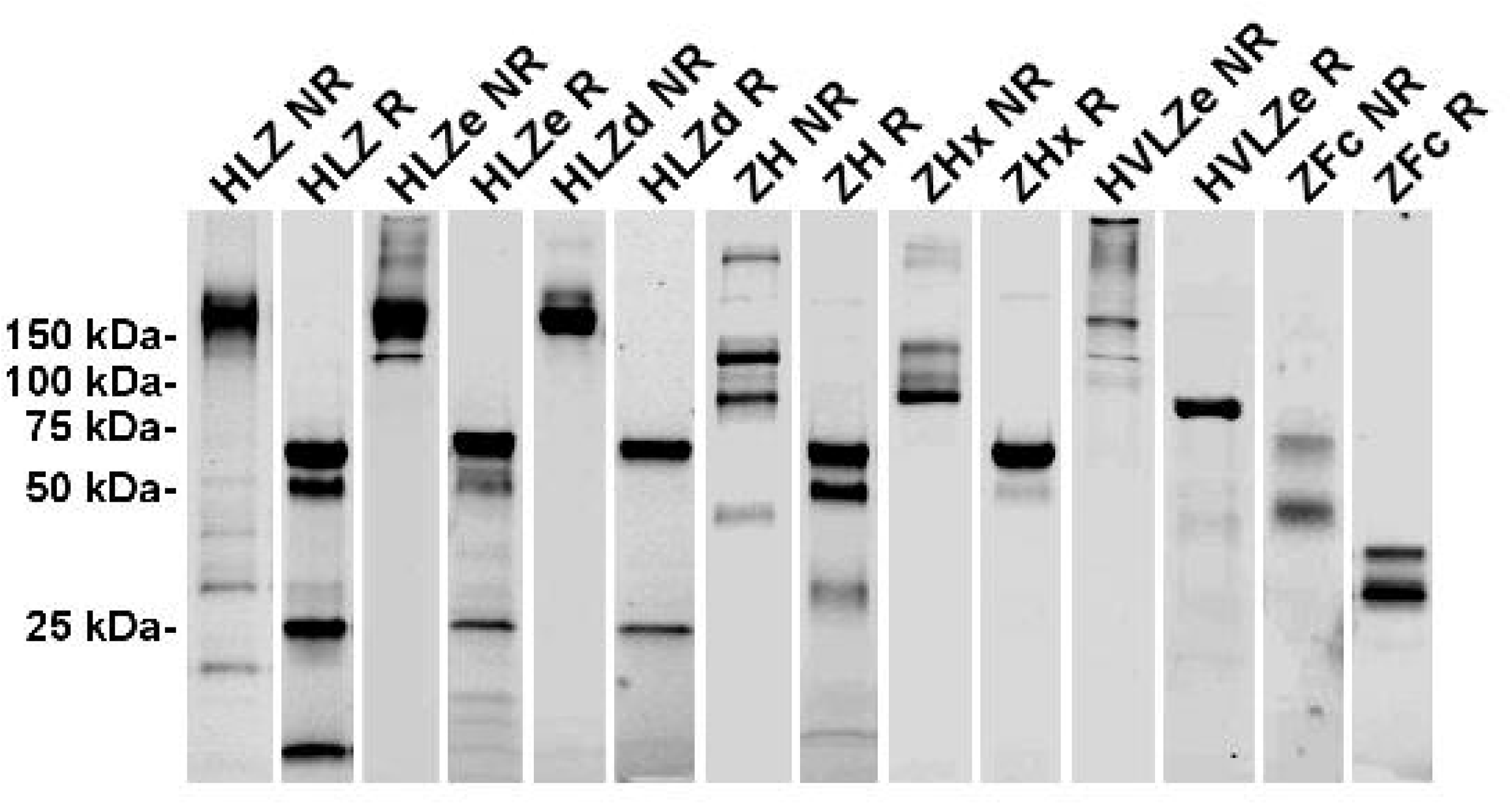
Purification of IgG Fusions. Agroinfiltrated leaf material from between 1-3 plants per construct was homogenized, clarified, and purified by protein G affinity chromatography. The peak elutions were pooled and separated on nonreducing (NR) and reducing (R) SDS-PAGE using stain-free polyacrylamide gels. Representative lanes for each construct are compiled from multiple gels here.

**Figure 4.**
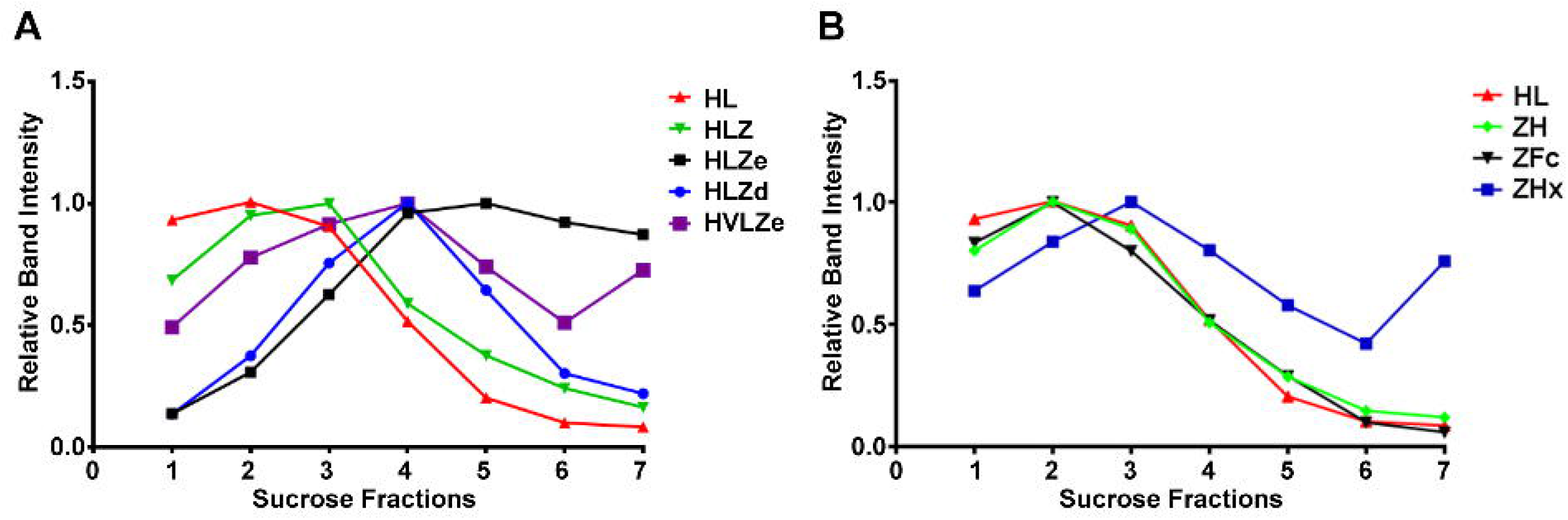
Sucrose gradient centrifugation of IgG fusions. Purified IgG fusions were separated by sucrose gradient sedimentation using 5/10/15/20/25% discontinuous sucrose layers. Gradient fractions were analyzed by SDS-PAGE and representative results are shown; direction of sedimentation is left to right. The relative band intensity was quantified using ImageJ software and the peak band was arbitrarily assigned the value of 1.

### 2.5 Polymeric constructs have improved stability

Producing vaccines which are stable at refrigerated temperatures, ambient temperatures, and after freeze/thawing, is a major challenge of the vaccine industry (Kumru et al., 2014). First, the stability of each construct was analyzed by comparing fully formed products and degradation products on SDS-PAGE gels after treatment with various temperature conditions. The initial level of degradation, which corresponds to any degradation that occurred during expression, purification, or the initial freeze/thaw, was substantially reduced (20-40% more fully formed product) in polymeric constructs when compared to monomeric constructs (**Fig. 5A**). After five freeze-thaw cycles or two weeks at 4°C, small amounts of degradation (2-5%) were observed with all constructs (**Fig. 5A**). High concentrations (>1 mg/ml) of HLZe and HVLZe precipitated after several days at 4°C (data not shown), probably due to the formation of very large insoluble complexes, thereby highlighting the undesirability of these constructs. At room temperature, most constructs had 10-15% degradation after two weeks (**Fig. 5A**). Overall, the polymeric constructs retained the highest stability, while the monomeric constructs, and especially the Fc fusion, degraded more rapidly (**Fig. 5A**).

**Figure 5.**
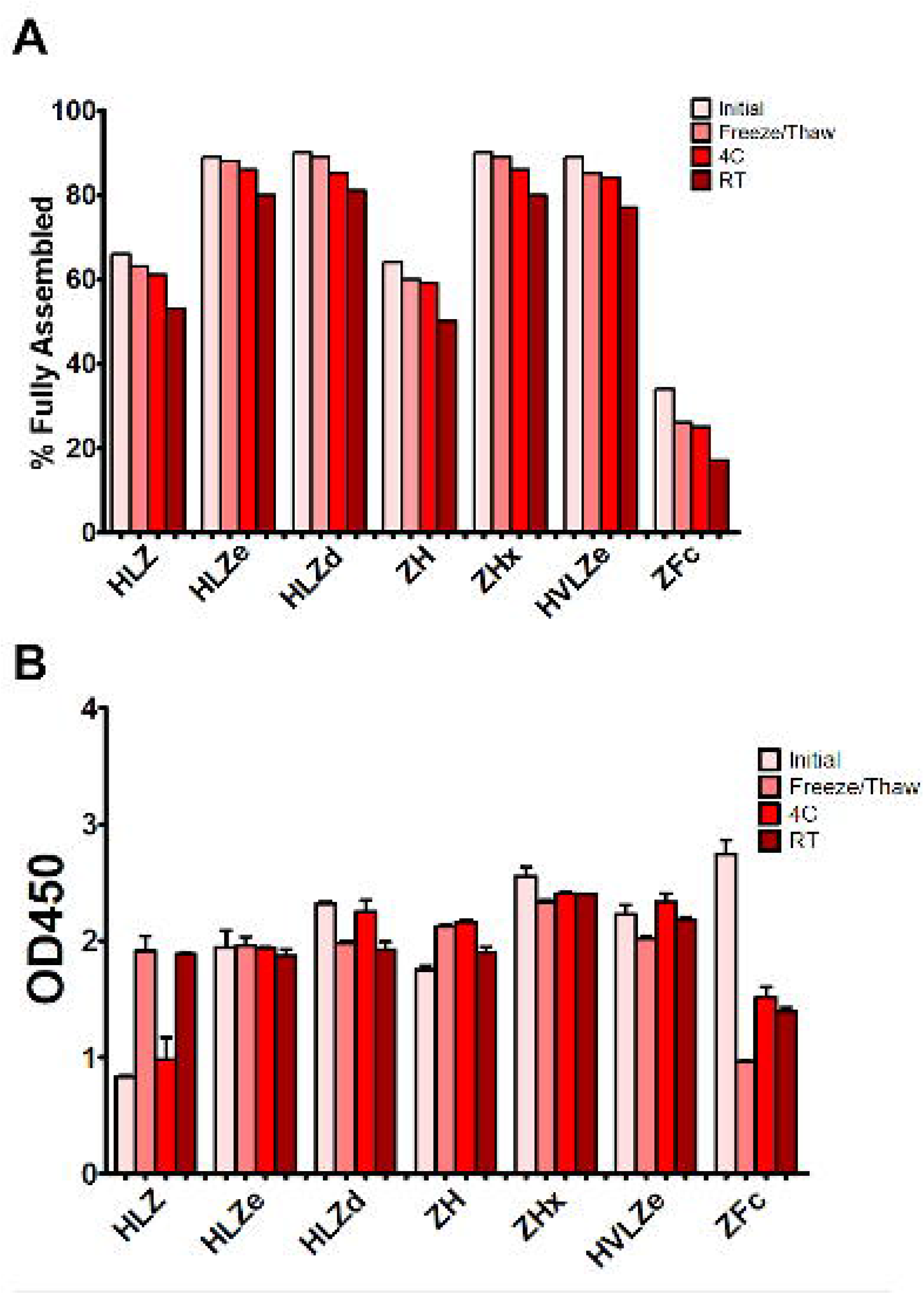
Stability and C1q binding of purified IgG fusions. Samples of purified IgG fusions were frozen and thawed once after purification (initial), or additionally subjected to either additional 5 freeze/thaw cycles, incubation for 2 weeks at 4°C, or incubation for 2 weeks at 23°C. (A) After each treatment, samples were separated on reducing and nonreducing SDS-PAGE gels, and the relative proportion of fully assembled product was analyzed using ImageJ software. The initial measurement reflects degradation which occurred during expression or purification, whereas the other measurements reflect degradation which occurred during each respective treatment. (B) Samples from each treatment were analyzed by C1q binding ELISA. Columns represent the mean OD_450_ value ± standard error from three samples.

Second, we determined whether storage resulted in an impairment of C1q binding. Most constructs retained strong C1q binding after storage; however, after room temperature storage, repeated freeze-thaw cycles, but not 4°C storage, construct HLZ unexpectedly displayed increased C1q binding (**Fig. 5B**). This may be due to aggregation, degradation of the Fab regions, or loss of light chain as numerous degradation products are visible on SDS-PAGE (**Fig. 3**). Conversely, construct ZFc lost C1q binding ability as it became more degraded, especially after freeze-thaw cycles, suggesting degradation of the Fc binding region (**Fig. 5B**). Taken together with the expression data, constructs with intermediate levels of self-binding (e.g. HLZd and ZHx) showed the highest stability during both extraction and storage while maintaining high solubility. Of these, only construct HLZd could be produced at high levels.

### 2.6 IgG fusion enhances ZE3 immunogenicity

IgG fusions have strong immunological properties even in the absence of adjuvant due to their improved binding of C1q and uptake via FcRn and Fc receptors on antigen presenting cells, (Czajkowsky et al., 2012). To investigate the immunogenicity of the IgG fusions created here, BALB/c mice (n = 6) were immunized subcutaneously, without adjuvant, with two doses of each IgG fusion construct such that each dose of ZE3 delivered was 8 μg. As a control, mice were also immunized with 8 μg His-tagged plant-expressed ZE3. As expected, all IgG fusions very strongly enhanced the production of ZE3-specific IgG, producing 20-fold to 150-fold higher total IgG titers than ZE3 alone (**Fig. 6A**, p < 0.01 compared to ZE3). 5-fold to 10-fold higher mean titers were reached by the ZHx and ZFc constructs compared to the other constructs, though these differences were not statistically significant. The level of IgG2a antibodies were also measured because they have important antiviral effector functions (Lu et al., 2018) and are correlated with T-cell activation (Huber et al., 2006) and complement activity (West et al., 2018). All IgG fusions significantly enhanced the production of IgG2a compared to ZE3 alone (**Fig. 6B**, p < 0.01 compared to ZE3). The construct HLZ, which displayed the lowest C1q binding (**Fig. 1B**), had a significantly reduced production of IgG2a compared to most other fusions (**Fig. 6B**, p < 0.05 compared to HLZ). Compared to total IgG, the relative ratio of IgG2a was lowest in the ZE3, HLZ, and ZFc groups (**Fig. 6B**, see table). Sera from mice immunized with all IgG fusions neutralized ZIKV (**Fig 7,**p < 0.0001 compared to sera from PBS-injected mice). All IgG fusions elicited significantly higher neutralizing responses than His-tagged ZE3 (**Fig 7**, p < 0.024 compared to ZE3).

**Figure 6.**
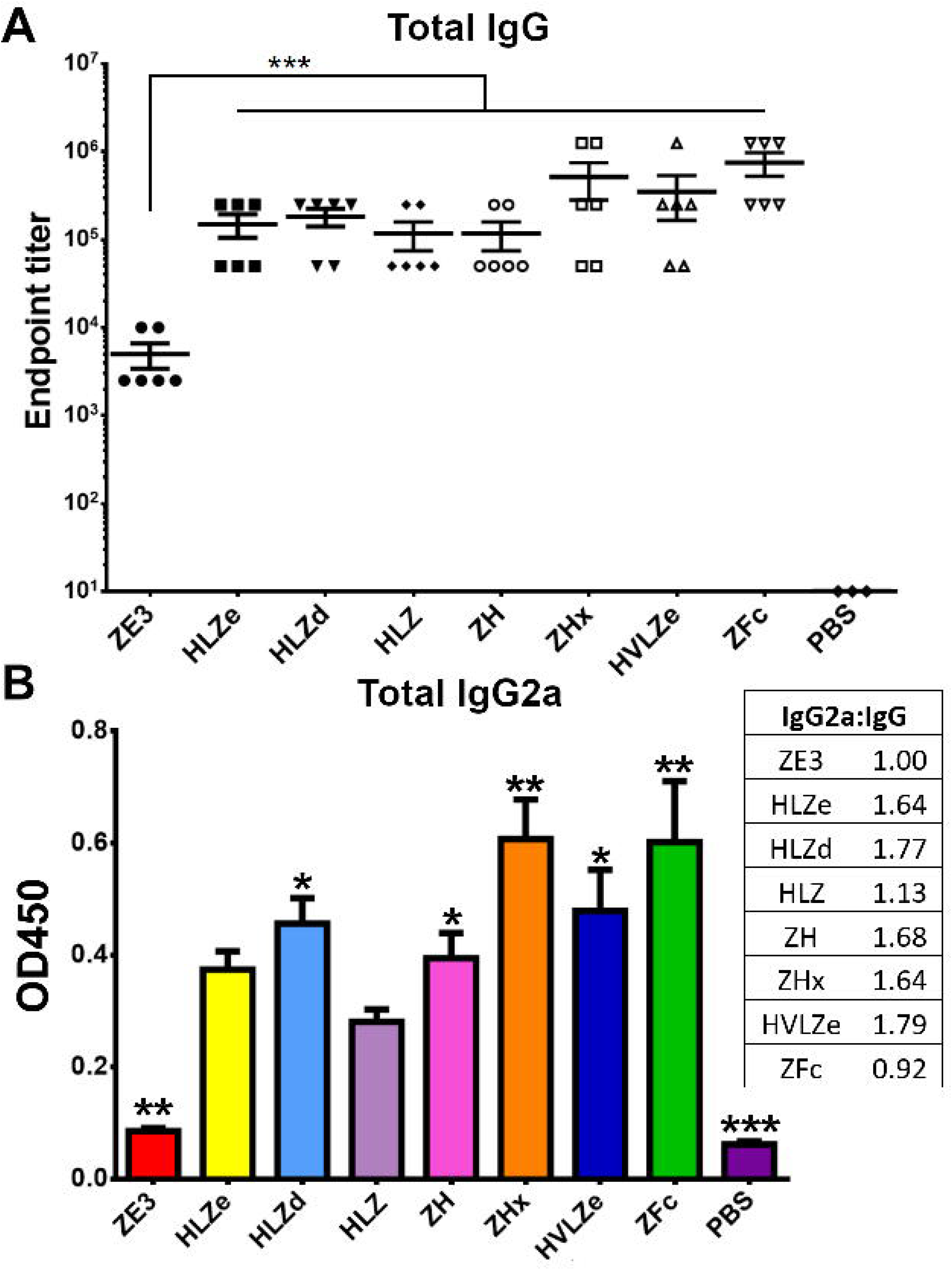
Mouse immunization and serum titers. BALB/c mice (6 per group) were immunized twice two weeks apart subcutaneously with a dose that would deliver 8 μg ZE3 for each IgG fusion or with PBS as a control. Mouse serum samples were collected two weeks after the final dose. (A) Serially diluted mouse serum was analyzed for total IgG production by ELISA. The endpoint was taken as the reciprocal of the greatest dilution that gave an OD_450_ reading at least twice the background. Two stars (**) indicates p < 0.01 by one-way ANOVA comparing the indicated columns to ZE3. (B) Mouse serum samples were diluted 1:100 and analyzed for IgG2a production by ELISA. The table shows the ratio of IgG2a to total IgG for each group, calculated as relative to ZE3, with the ZE3 value set arbitrarily at 1.0. Values with one star (*) indicates p < 0.05 and (**) indicates p < 0.01 by one-way ANOVA comparing the indicated columns to HLZ.

**Figure 7.**
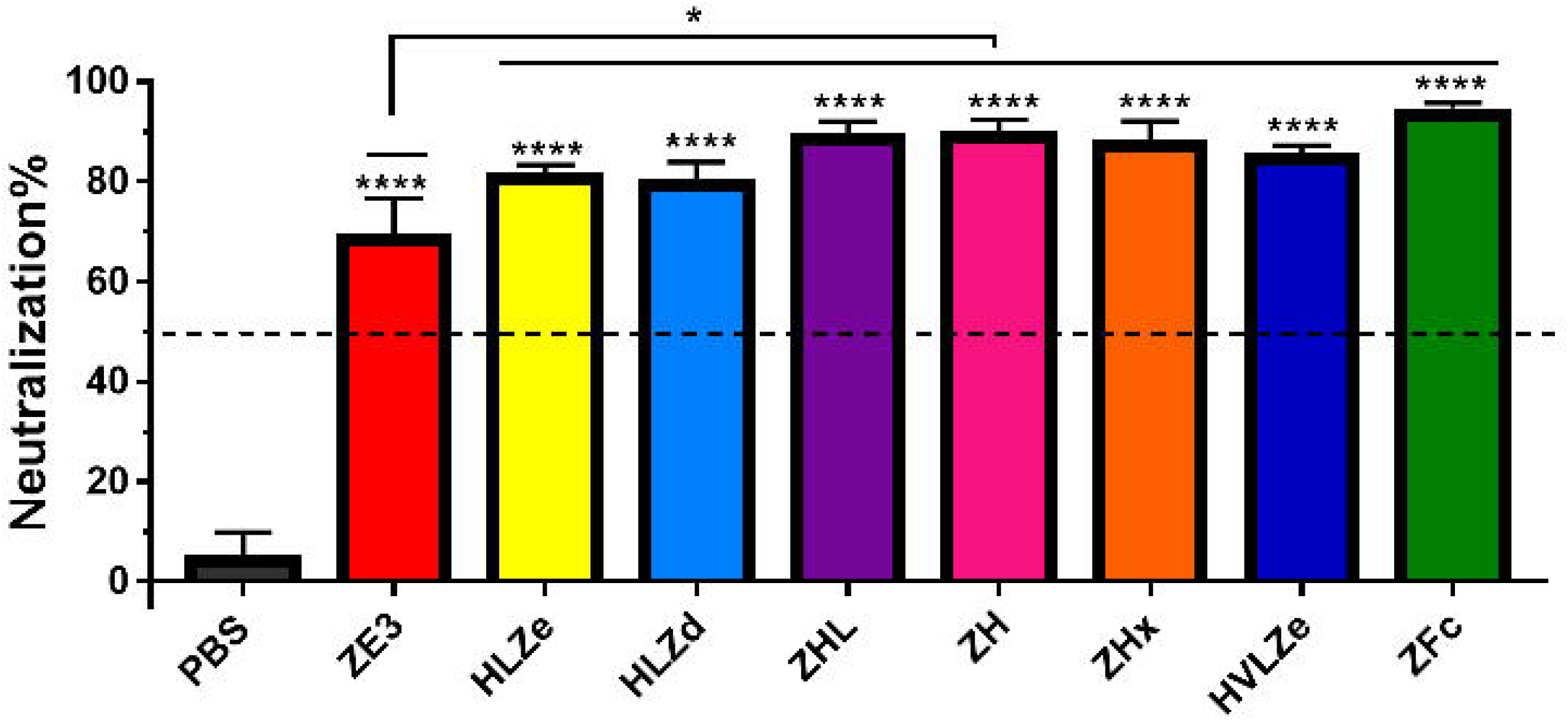
Neutralization of ZIKV. Plaque reduction neutralization assays (PRNT) assays were performed using pooled mouse sera with dilution ratio of 1:10 to evaluate ZIKV-specific neutralizing antibodies in the sera. Data are presented as mean neutralization % and SD from three independent experiments with technical triplicates for each sample. Statistical analyses were performed using one-way ANOVA, p values from comparison between vaccine treatments and PBS were indicated with **** (<0.0001) or from comparison between vaccine treatments and His-tagged ZE3 were indicated with * (p < 0.024).

## 3. Discussion

Previous work has shown that IgG fusion strategies have the potential to enhance antigen immunogenicity (Kim et al., 2018; Konduru et al., 2011; Loureiro et al., 2011; Webster et al., 2018; Zhao et al., 2018). However, the key characteristics to produce an optimal IgG fusion vaccine candidate have not been directly compared. An optimal vaccine candidate must be accessible, affordable, and stable, as well as immunogenic. Impressively, especially given the low doses and lack of adjuvant used in this study, the IgG fusions were potently immunogenic with many of the best candidates eliciting 100-fold higher antibody responses compared to antigen alone. While the highest immunological readouts were obtained by constructs ZHx and ZFc, these constructs have limited use as vaccine candidates, as ZHx suffered from low yield and aggregation, while ZFc was extraordinarily unstable prior to and during storage. These findings underscore the importance of evaluating multiple aspects of vaccine viability. By contrast, construct HLZd had nearly double the yield of the next closest construct (**Fig. 2**) while maintaining high stability (**Fig. 5**), solubility (**Fig. S5B**), and immunological properties (**Figs. 1B, 6, 7**). All of the HLZd produced in this study was made in a single small *N. benthamiana* plant (data not shown), demonstrating the excellent potential of plant-based vaccine development to address vaccine affordability and accessibility. The overall properties of each construct are summarized in Table 1. These findings warrant further development of HLZd-like constructs as a vaccine platform which we anticipate will be broadly applicable to the development of vaccines targeting other pathogens.

**Table 1.** Summary of characteristics of IgG fusions. A summary of the characteristics of each IgG fusion vaccine candidate is given. For expression, only the yield (mg/g LFW) of the fully assembled product is shown. A greater number of “+” symbols indicates either a statistically significant increase in the mean value for that property (for C1q, IgG, IgG2a, and neutralization), or a repeatably observed difference (for sedimentation). For ZHx (*), peaks of both low-and high-velocity sedimentation were observed.

| Construct | Yield | Solubility | Sedimentation | Stability | C1q | IgG | IgG2a | Neutralization |
| --- | --- | --- | --- | --- | --- | --- | --- | --- |
| ZE3 | N.D. | N.D. | N.D. | N.D. | N.D | + | + | ++ |
| HLZe | 0.22 mg/g | Low | +++++ | High | +++ | +++ | ++ | +++ |
| HLZd | 1.50 mg/g | High | +++ | High | +++ | +++ | +++ | +++ |
| HLZ | 0.17 mg/g | High | ++ | Medium | + | +++ | ++ | +++ |
| ZH | 0.83 mg/g | High | + | Medium | ++ | +++ | +++ | +++ |
| ZHx | 0.21 mg/g | Low | +++* | High | +++ | +++ | ++++ | +++ |
| VHLZe | <0.1 mg/g | Low | +++++ | High | +++ | +++ | +++ | +++ |
| ZFc | 0.57 mg/g | High | + | Low | +++ | +++ | ++++ | +++ |

While most current antibody fusion vaccines use constructs similar to ZFc, many properties of Fc fusions, including C1q binding, vary based on the individual fusion partner and thus must be determined empirically for each fusion (Lagassé et al., 2019). Interestingly, some Fc fusion constructs form hexamers (Zhang et al., 2019), which may explain the strong C1q binding and immunogenicity of ZFc observed here. Despite its strong immunogenicity, ZFc was remarkably unstable: ~50% of the ZE3 was proteolytically cleaved either before or during extraction (**Fig. 2A, 2B**), even when directly extracted in SDS sample buffer or with protease inhibitors (data not shown). Further, less than 25% of full-size ZFc molecules remained intact upon repeated freeze-thaw or storage (**Fig. 5A**). This degradation was associated with a loss in C1q binding (**Fig. 5B**), probably due to impairment of the Fc receptor binding domains. ZFc has fewer disulfide bonds than whole IgG, and in general Fc fusions often suffer from instability or undesirable aggregation (Zeng et al., 2018). Further protein engineering, including optimization of linkers, may improve the stability and expression of these constructs. In contrast, IgG fusions displaying self-binding properties, such as HLZd, HLZe, and ZHx, were much more stable during extraction and storage. Consistent with their reduced binding to antibody probes during ELISA (**Fig. 2A**), the intramolecular and intermolecular associations of these constructs may protect them from proteolysis or other degradative processes.

The large complex size generated by traditional RIC (i.e. HLZe) renders them poorly soluble upon extraction (**Figs. 2A/S2,**Diamos et al., 2019) with concentrations above 1-2 mg/ml precipitating during storage (data not shown). Therefore, we reduced the binding affinity of the RIC by mutating the 6D8 epitope tag (construct HLZd). Using this strategy of modifying the strength of self-binding, RIC can be customized to create complexes of different sizes. These findings will allow future investigation of the immunological and functional properties conferred by variably sized immune complexes. While traditional RIC require co-expression of both the heavy and light chains, this can be simplified by creating a single chain antibody with the variable light domain fused to the variable heavy domain, as in construct HVLZe. This construct had high stability (**Fig. 5**) and immunogenicity (**Fig. 6**), but was poorly soluble, resulting in low recovery (**Fig. 2**). By further reducing the epitope binding of HVLZe or by altering the relative positions of the light chain or antigen fusions, or by using the shorter “d” epitope tag, it is likely the solubility can be improved.

The glycosylation state of the Fc strongly controls its function and can enhance or inhibit binding to immune receptors by modulating the stability, conformation, and aggregation of the Fc (Mastrangeli et al., 2019). These alterations result in important differences in antibody effector functions, including antibody-dependent cellular cytotoxicity, antibody-dependent cell-mediated phagocytosis, complement-dependent cytotoxicity, and antibody-dependent enhancement of viral infection (Sun et al., 2018). Advances in glycoengineering have allowed targeted optimization of the Fc glycosylation state in a variety of recombinant expression systems (Gupta and Shukla, 2018; Kallolimath and Steinkellner, 2015; Wang et al., 2018). For example, an anti-CD20 antibody produced in glycoengineered plants lacking core xylose and fucose N-linked glycans showed improved binding to FcγRI, FcγRIIIa, and C1q (Marusic et al., 2018). Similarly, anti-DENV antibodies produced in glycoengineered plants have been shown to have diminished ADE activity and, consequently, have superior efficacy and safety profiles than their mammalian cell-produced counterparts (Dent et al., 2016). Antibody therapeutics made in glycoengineered plants have also been used to successfully treat rhesus macaques and humans with Ebola (Lyon et al., 2014; Olinger et al., 2012; Qiu et al., 2014; Zeitlin et al., 2011), HIV (Forthal et al., 2010; Ma et al., 2015; Stelter et al., 2020), and Chikungunya virus disease (Hurtado et al., 2019). In agreement with these studies, we found that C1q binding was enhanced in glycoengineered plants in a manner which acted synergystically with immune complex formation (**Fig. S1A**). As the entire mamallian sialylation pathway has been reconsistuted in glycoengineered plants (Castilho et al., 2012), and sialylated immune complexes yield higher affinity antibodies (Maamary et al., 2017), the IgG fusion vaccines used in this study may be improved by further glycan modification.

Despite its theoretically monomeric nature, construct HLZ elicited high titers and comparable neutralization to the more polymeric constructs. HLZ produced a small but repeatable improvement in C1q binding (**Fig. 1B**) and a shift in small shift in density (**Fig. 4A**, compare HL and HLZ) which may suggest some low-level aggregation due to the ZE3 fusion. Nonetheless, the expression of HLZ was generally low, suggesting C-terminal ZE3 fusion may interfere with proper folding or otherwise cause instability (**Fig. 2A**). Many degradation products are visible by SDS-PAGE following purification (**Fig. 3**) and, interestingly, the C1q binding of HLZ continued to increase upon further degradation by incubation for 2 weeks at RT or repeated freeze-thaw cycles (**Fig. 5B**). These data suggest that the degradation products may be more highly immunogenic than the original construct, as light chain or Fab removal strongly improves C1q binding (**Fig. S2**). This finding may have also contributed to the relatively high immunogenicity produced by HLZ (**Fig. 6A**). However, despite similar total IgG titers and ZIKV neutralization, the relative IgG2a ratio was significantly reduced for HLZ, ZE3, and ZFc when compared to the more polymeric constructs (**Fig. 6B**). Since IgG2a antibodies are a general indicator of a Th1 response (Huber et al., 2006) and complement activation is involved in T-cell immunity (West et al., 2018), these constructs may have reduced T-cell activation compared to more polymeric constructs. This is important because T-cell activation and complement activation are vital components to generate long-term B-cell immunity (Akkaya et al., 2020; West et al., 2018). *In vitro* neutralization (**Fig. 7**) does not account for antibody subclass effector functions (e.g. IgG2a) or cellular immunity, thus further investigations are warranted.

Another point to consider is that, at least for certain antibody/antigen combinations, antigen binding induces conformational changes in IgG, resulting in improved hexamer formation and subsequently improved C1q binding (Wang et al., 2016). Mixing 6D8 with an antigen containing the 6D8 epitope produced only a small increase in C1q binding (**Fig. S2**). However, antigen fusion to the 6D8 N-terminus had a more pronounced effect on C1q binding (**Fig. 1**), perhaps by strongly inducing conformational changes similar to antigen binding which allow hexamer formation (Diebolder et al., 2014). In general, we have found that antigen fusion to the 6D8 N-terminus greatly enhances C1q binding for a variety of large and small antigens (data not shown). Furthermore, removal of the 6D8 light chain (construct ZH) also substantially enhanced C1q binding (**Fig. 1**). These findings agree with the hypothesis that the Fab portions of the antibody plays a regulatory role in C1q binding. Additionally, removal of the light chain may also impact glycosylation and thus C1q binding (He et al., 2014).

In summary, we have developed a cheap, fast, and efficient plant expression system to produce and purify high levels of IgG fusion vaccine candidates. Plant recombinant expression systems are particularly well suited to make IgG fusion vaccines, such as the ones described in this study, since they have inherent safety, high scalability, and low production costs when compared to mammalian cell systems (Alam et al., 2018; Buyel, 2019; Chen and Davis, 2016; Gleba et al., 2014). These benefits can be of special value for vaccine production in developing countries (Ma et al., 2013). By directly comparing many different IgG fusion strategies, we have identified a self-assembling immune complex construct HLZd that is highly expressing, stable, soluble, and immunogenic. These studies warrant future research in animal models, and we anticipate our findings will be broadly applicable to other vaccine antigens or antibody-based therapeutics.

## 4. Experimental Procedures

### 4.1 Vector construction

The construction of a geminiviral replicon plant expression vector for ZE3, as well as its fusion to the 6D8 C-terminus (pBYR11eM-h6D8ZE3, referred to here as construct “HLZe”) or N-terminus with epitope tag (pBYR11eMa-BAZE3-Hgp371) or without epitope tag (pBYR11eMa-BAZE3-H) have been previously described (Diamos et al., 2020b, 2020a) A vector pBYKEMd2-6D8 expressing the full 6D8 mAb without ZE3 fusion (construct “HL”) has been previously described (Diamos et al., 2020b, 2020a). To create a vector expressing only the light chain of 6D8, pBYKEMd2-6D8 was digested with XhoI and the vector was self-ligated to yield pBYKEMd-6D8K. A vector expressing only the heavy chain of 6D8 (construct “H”) was created by digesting pBYKEMd2-6D8 with SacI and self-ligating the vector, to yield pBYKEMd-6D8H. The 6D8 epitope binding tag was added to pBYKEMd-6D8H by digesting pBYR11eMa-BAZE3-Hgp371 with BsaI-SacI and inserting the tag-containing fragment into pBYKEMd-6D8H digested with BsaI-SacI, yielding pBYKEMd-6D8Hgp371 (construct “HLe” when coexpressed with light chain). To remove the epitope tag from HLZe, pBYR11eM-h6D8ZE3 was digested with BamHI-SacI and ligated with a fragment containing ZE3 obtained via amplification with primers ZE3-Bam-F (5’-gcgggatccaagggcgtgtcatactcc) and ZE3-Sac-R (5’-acagagctcttaagtgctaccactcctgtg) and subsequent digestion with BamHI-SacI. The resulting vector, pBYKEMd-HZE3, was coinfiltrated with pBYKEMd-6D8K to produce construct “HLZ.” To produce ZE3 fused to the 6D8 N-terminus without light chain, pBYR11eMa-BAZE3-H was digested with SacI and the vector vector was self-ligated, yielding pBYKEMd-ZE3H (construct “ZH”). To introduce hexamer mutations, a region of the 6D8 heavy chain constant region was synthesized (Integrated DNA Technologies, Iowa, USA) containing the E345R, E430G, and S440Y mutations, then digested with BsaI-SacI and used to replace the BsaI-SacI region of 6D8 in pBYKEMd-ZE3H, yielding pBYKEMd-ZE3Hx (construct “ZHx”). RIC epitope tag mutant “a” was generated by annealing oligos 6D89-F (5’-ctagtgtttacaagctggacatatctgaggcataagagct) and 6D89-R (5’-cttatgcctcagatatgtccagcttgtaaaca) and ligating them into pBYR11eM-h6D8ZE3 digested SpeI-SacI; mutant “b” was generated by first amplifying mutant “a” with primers gpDISE-Sac-R: (5'-tttgagctcttactcagatatgtccagcttgtaaac) and 35S-F (5’aatcccactatccttcgc), then digesting the product with SpeI-SacI and ligating it into pBYR11eM-h6D8ZE3 digested with SpeI-SacI. Mutants “c” and “d” were created similarly to mutant “a” using overlapping oligos 6D87-F (5'-ctagttacaagctggacatatctgagtaagagct) and 6D87-R (5’-cttactcagatatgtccagcttgtaa) for “c” and 6D86-F (5’-ctagttacaagctggacatatcttaagagct) and 6D86-R (5’-cttaagatatgtccagcttgtaa).

In order to make a construct in which the variable heavy (VH) domain is linked to a variable light chain (VL) domain that, in turn, is directly fused to the constant region of the 6D8 antibody, the variable regions were first obtained through PCR amplification and end-tailoring of segments from pBYR11eM-h6D8ZE3. For the VH domain, the primers LIR-H3A (5’-aagcttgttgttgtgactccgag) and 6D8VH-Spe-R (5’-cggactagtagctgaagacactgtgac) were used. The VL region was obtained through PCR amplification of pBYR11eM-h6D8ZE3 with primers 35S-F (5’-aatcccactatccttcgc) and 6D8VK-Nhe-R (5’-cgtgctagccttgatctccactttggtc). In order to fuse VL region to the constant region of a human IgG antibody, a subclone was created by digesting the PCR fragment with XhoI-NheI and inserting it into a vector, pKS-HH-gp371, that contained the 6D8 heavy chain (Kim et al., 2015). This subclone was named pKS-VL. Next, pBYKEM-6D8K was digested with SbfI-SacI, the PCR product that amplified the variable heavy chain fragment was digested SbfI-SpeI, and the variable light chain subclone was digested SpeI-SacI. These fragments were assembled to create pBYKEMd2-VHLVK (construct “HVL”). Finally, this construct was used to create pBYKEMd2-HVLZe by a two-fragment ligation. The pBYKEMd2-VHLVK construct was digested BsaI and SacI to obtain the vector fragment along with the variable regions of the heavy and light chains. To obtain the ZE3 antigen segment and the epitope tag, pBYR11eM-h6D8ZE3 was also digested BsaI-SacI. The resulting construct, which was used to produce HVLZe, was named pBYKEMd-HVLZe.

### 4.2 Agroinfiltration of *Nicotiana benthamiana* leaves

Binary vectors were separately introduced into *Agrobacterium tumefaciens* EHA105 by electroporation. The resulting strains were verified by restriction digestion or PCR, grown overnight at 30°C, and used to infiltrate leaves of 5-to 6-week-old *N. benthamiana* maintained at 23-25°C. Briefly, the bacteria were pelleted by centrifugation for 5 min at 5,000g and then resuspended in infiltration buffer (10 mM 2-(N-morpholino)ethanesulfonic acid (MES), pH 5.5 and 10 mM MgSO4) to OD600=0.2, unless otherwise described. The resulting bacterial suspensions were injected by using a syringe without needle into leaves through a small puncture (Huang and Mason, 2004). To evaluate the effects of glycosylation, transgenic plants silenced for xylosyltransferase and fucosyltransferase were employed (Castilho and Steinkellner, 2012). Plant tissue was harvested at 5 days post infiltration (DPI).

### 4.3 Protein extraction, expression and purification

Crude protein was extracted by homogenizing agroinfiltrated leaf samples with 1:5 (*w:v*) ice cold extraction buffer (25mM Tris-HCl, pH 8.0, 125mM NaCl, 3mM EDTA, 0.1% Triton X-100, 10 mg/mL sodium ascorbate, 0.3 mg/mL phenylmethylsulfonyl fluoride) using a Bullet Blender machine (Next Advance, Averill Park, NY) following the manufacturer’s instruction. Homogenized tissue was rotated at 4°C for 30 min. The crude plant extract was clarified by centrifugation at 13,000*g* for 15 min at 4°C and the supernatant was analyzed by SDS-PAGE or ELISA. Alternatively, to evaluate solubility of proteins in the original homogenate, the pellet was designated the insoluble fraction and treated with SDS sample buffer at 100°C for 10 min before loading on SDS-PAGE.

IgG variants were purified by protein G affinity chromatography. Agroinfiltrated leaves were blended with 1:3 (w:v) ice cold extraction buffer (25mM Tris-HCl, pH 8.0, 125mM NaCl, 3mM EDTA, 0.1% Triton X-100, 10 mg/mL sodium ascorbate, 0.3 mg/mL phenylmethylsulfonyl fluoride), stirred for 30 min at 4°C, and filtered through miracloth. To precipitate endogenous plant proteins, the pH was lowered to 4.5 with 1M phosphoric acid for 5 min while stirring, then raised to 7.6 with 2M Tris base. Following centrifugation for 20 min at 16,000*g*, the clarified extract was loaded onto a Protein G column (Thermo Fisher Scientific, Waltham, MA, USA) following the manufacturer’s instructions. Purified proteins were eluted with 100mM glycine, pH 2.5, directly into collection tubes containing 1M Tris-HCl pH 8.0 to neutralize the elution buffer and stored at −80°C. Purified protein concentration was measured by A280 absorbance, ELISA, and gel quantification.

ZE3-His expressed from pBYe3R2K2Mc-BAZE3 was purified by metal affinity chromatography. Protein was extracted as described above, but without acid precipitation. The clarified extract was loaded onto a column containing TALON Metal Affinity Resin (BD Clontech, Mountain View, CA) according to the manufacturer’s instructions. The column was washed with PBS and eluted with elution buffer (PBS, 150mM imidazole, pH 7.4). Peak ZE3 elutions were pooled, dialyzed against PBS, and stored at −80°C.

### 4.5 SDS-PAGE and Western blot

Plant protein extracts or purified protein samples were mixed with SDS sample buffer (50 mM Tris-HCl, pH 6.8, 2% SDS, 10% glycerol, 0.02 % bromophenol blue) and separated on 4-15% stain-free polyacrylamide gels (Bio-Rad, Hercules, CA, USA). For reducing conditions, 0.5M DTT was added, and the samples were boiled for 10 min prior to loading. Polyacrylamide gels were visualized and imaged under UV light, then transferred to a PVDF membrane. For IgG detection, the protein transferred membranes were blocked with 5% dry milk in PBST (PBS with 0.05% tween-20) overnight at 4°C and probed with goat anti-human IgG-HRP (Sigma-Aldrich, St. Louis, MO, USA diluted 1:5000 in 1% PBSTM). Bound antibody was detected with ECL reagent (Amersham, Little Chalfont, United Kingdom).

### 4.6 ELISA quantification of construct expression levels

For ZE3 ELISA, a 96-well high-binding polystyrene plate (Corning Inc, Corning, NY, USA) was coated with a 1:500 dilution of unlabeled goat anti-human IgG (Southern Biotech, Birmingham, AL, USA) and incubated at 37°C for 1 h. After being washed with PBST, the plates were blocked with 5% non-fat dry milk in PBST for 30 min and washed with PBST. Multiple dilutions, ranging from 1:100 to 1:25,000, of clarified plant extracts containing the different constructs were added to the plate. Previously purified and quantified plant-produced HLZ was included as a standard and positive control while samples of uninfiltrated crude extract were added for a negative control. After a 1-h incubation, the plate was washed three times in 1x PBST. Plates were then incubated with a 1:1000 dilution of mouse serum raised against ZE3-6H followed by goat anti-mouse IgG HRP conjugate (Southern Biotech, Birmingham, AL, USA) for 1 h at 37°C. The plate was then washed five times with PBST, developed with TMB substrate (Thermo Fisher Scientific, Waltham, MA, USA), and the absorbance read at 450 nm. IgG ELISA was performed similarly, except constructs were bound directly to the plate and detected with goat anti-mouse IgG HRP conjugate (Southern Biotech, Birmingham, AL, USA).

### 4.6 C1q binding

96-well high-binding polystyrene plates (Corning Inc, Corning, NY, USA) were coated with 15 μg/ml human complement C1q (PFA, MilliporeSigma, MA) in PBS for 2h at 37°C. The plates were washed 3 times with PBST, and then blocked with 5% dry milk in PBST for 15 minutes. After washing 3 times with PBST, purified human IgG (Southern Biotech, Birmingham, AL, USA) and purified IgG-ZE3 fusions were added at 0.1 mg/ml with 10-fold serial dilutions and incubated for 1.5 hours at 37°C. After washing 3 times with PBST, bound IgG was detected by incubating with 1:1000 polyclonal goat anti human IgG-HRP (Southern Biotech, Birmingham, AL, USA) for 1h at 37°C. The plates were washed 4 times with PBST, developed with TMB substrate (Thermo Fisher Scientific, Waltham, MA, USA), stopped with 1M HCl, and the absorbance was read at 450nm.

### 4.7 6D8 epitope binding

To test the ability of HVL to bind to the 6D8 epitope tag, 900 ng of purified dengue consensus envelope domain III tagged with 6D8 epitope (Kim et al., 2015) were bound to a 96-well high-binding polystyrene plate (Corning Inc, Corning, NY, USA). After a 1-hour incubation at 37°C, the plate was washed thrice with PBST and blocked with 5% dry milk in PBST for 30 minutes. Then, the plate was washed thrice with PBST and various dilutions of either purified HLV or full-length 6D8 antibody were added to the plate. The plate was incubated at 37°C for 1-hour, washed thrice with PBST and detected with HRP-conjugated mouse anti-human IgG (Fc only) (Southern Biotech, Birmingham, AL, USA) antibody at a 1:2000 dilution. Then, the plate was thoroughly washed with PBST and developed with TMB substrate (Thermo Fisher Scientific, Waltham, MA, USA). The absorbance was read at 450nm.

### 4.8 Sucrose gradient density centrifugation

Purified samples of each IgG fusion (100 µl) were loaded onto discontinuous sucrose gradients consisting of 350 µl layers of 5, 10, 15, 20, and 25% sucrose in PBS in a 2.0 ml microcentrifuge tubes and centrifuged at 21,000*g* for 16 h at 4 °C. Fractions were collected from the top and analyzed by SDS-PAGE, followed by visualization on stain-free gels (Bio-Rad, Hercules, CA, USA). The relative band intensity of each fraction was determined using ImageJ software, with the peak band arbitrarily assigned the value of 1.

### 4.9 Stability of IgG fusions during storage

After purification, samples of purified IgG fusions were frozen at −80°C. Samples were either untreated (referred to as “initial”), or thawed and then subjected to either five additional freeze/thaw cycles, or incubated for 2 weeks at 4°C, or incubated for 2 weeks at 23°C. Initial and treated samples were visually inspected for any signs of precipitation, and then analyzed by reducing and non-reducing SDS-PAGE to observe cleavage of ZE3 or other degradation products. ImageJ analysis was used to compare the band intensity of the fully formed product to that of any degradation products. Each sample was also analyzed as described in section 4.6 for C1q binding.

### 4.10 Immunization of mice and sample collection

All animals were handled in accordance to the Animal Welfare Act and Arizona State University IACUC under study number 18-1616R. Female BALB/C mice, 6-8 weeks old, were immunized subcutaneously with purified IgG fusion variants. In all treatment groups, the total weight of antigen was set to deliver an equivalent 8 µg of ZE3. Doses were given on days 0 and 14. Serum collection was done as described (Santi et al., 2008) by submandibular bleed on days 0, 14, and 28.

### 4.11 Antibody measurements

Mouse antibody titers were measured by ELISA. Plant-expressed 6-His tagged ZE3 at 50 ng/well was bound to 96-well high-binding polystyrene plates (Corning Inc, Corning, NY, USA), and the plates were blocked with 5% nonfat dry milk in PBST. After washing the wells with PBST (PBS with 0.05% Tween 20), the mouse sera were diluted with 1% PBSTM (PBST with 1% nonfat dry milk) and incubated. Mouse antibodies were detected by incubation with polyclonal goat anti-mouse IgG-horseradish peroxidase conjugate (Sigma-Aldrich, St. Louis, MO, USA). The plate was developed with TMB substrate (Thermo Fisher Scientific, Waltham, MA, USA), stopped with 1M HCl, and the absorbance was read at 450nm. Endpoint titers were taken as the reciprocal of the lowest dilution which produced an OD_450_ reading twice the background produced using PBS as the sample. IgG2a antibodies were measured from sera diluted 1:100 in 1% PBSTM and detected with IgG2a horseradish peroxidase conjugate (Santa Cruz Biotechnology, Dallas, TX, USA). IgG2a values were normalized by total IgG, and the value obtained for ZE3 was arbitrarily defined as 1.0. The ratios for each construct were calculated as IgG2a:IgG values relative to that of ZE3.

### 4.12 Plaque reduction neutralization assay

Serum samples from terminal blood collection were pooled for each mouse group and heat inactivated. As described previously (Dent et al., 2016), the PRNT assay was carried out using mouse sera diluted in Opti-Mem media (Invitrogen) at ratio of 1:10. Each sample was incubated with 100 pfu ZIKV (PRVABC59, ATCC# VR-1843) in equal volume for 1hr at 37°C before the virus/serum mixture was added to each well of VERO cells (ATCC # CCL-81) in a 24 well plate. The virus/serum mixture was aspirated after a 1.5 hour incubation at 37°C. Subsequently, VERO cells were overlaid with 0.8% agarose in DMEM medium containing 5% FBS (Invitrogen, CA) and incubated at 37°C for three days. Finally, VERO cells were fixed with 4% paraformaldehyde (PFA, MilliporeSigma, MA) overnight and stained with 0.2% crystal violet. Plaques from each well were counted and neutralization % was calculated by using the formula: [(number of ZIKV plaque per well in virus only control wells)-(number of ZIKV plaque per well of diluted serum) / (number of ZIKV plaque per well in virus only control wells) × 100]. Neutralizing titers >10 refer to constructs with >50% neutralization at a 1:10 serum dilution.

## Abbreviations

RIC: recombinant immune complex
ZIKV: Zika virus
ZE3: Zika virus envelope domain III
Fc: the C-terminal fragment of crystallization of IgG
Fcγ: Fc from immunoglobulin G
Fab: the antigen-binding fragment of IgG
6D8: a human IgG1 monoclonal antibody targeting a linear epitope on Ebola virus glycoprotein 1
C1q: the first component of the classical complement pathway
ADCC: antibody-dependent cellular cytotoxicity
CDC: complement dependent cytotoxicity
FcRn: neonatal Fc receptors.

## Acknowledgements

This work was supported by funds provided by the School of Life Sciences, the Graduate and Professional Student Association, and the Center for Immunotherapy, Vaccines, and Virotherapy of the Biodesign Institute at Arizona State University. This work was also supported in part by a grant from National Institute of Allergy and Infectious Diseases (NIAID) # R33AI101329 to QC.

## Author Contributions

AD and HM designed experiments and analyzed data. AD, HM, and MP constructed vectors. AD performed C1q binding, solubility, sucrose gradient, and expression experiments. AD, JH, and MP performed purification experiments. AD and JH performed stability experiments. JK performed mouse immunization and bleeds. AD and MP performed antibody titer experiments. HS performed ZIKV neutralization experiments. AD wrote the manuscript. AD, HM, and QC critically revised the MS.

## Supporting Info

**Figure S1. C1q binding comparison between wildtype and glycoengineered plants**

**Figure S2. C1q binding comparison between IgG fusions**

**Figure S3. Epitope binding of single chain 6D8**

**Figure S4. RIC insolubility**

**Figure S5. Solubility and binding of 6D8 epitope tag mutants Tables**

