## Supplementary figures and images for "A highly expressing, soluble, and stable plant-made IgG fusion vaccine strategy enhances antigen immunogenicity in mice without adjuvant"

### Supplemental Figure 1

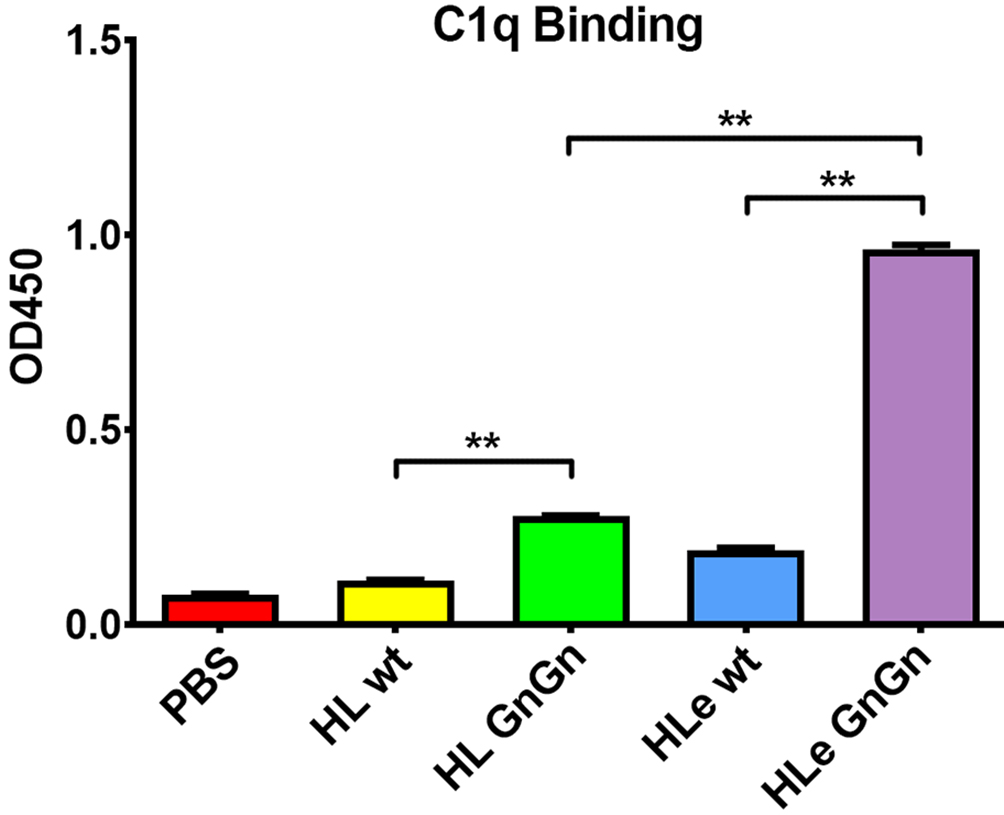

### Supplemental Figure 2

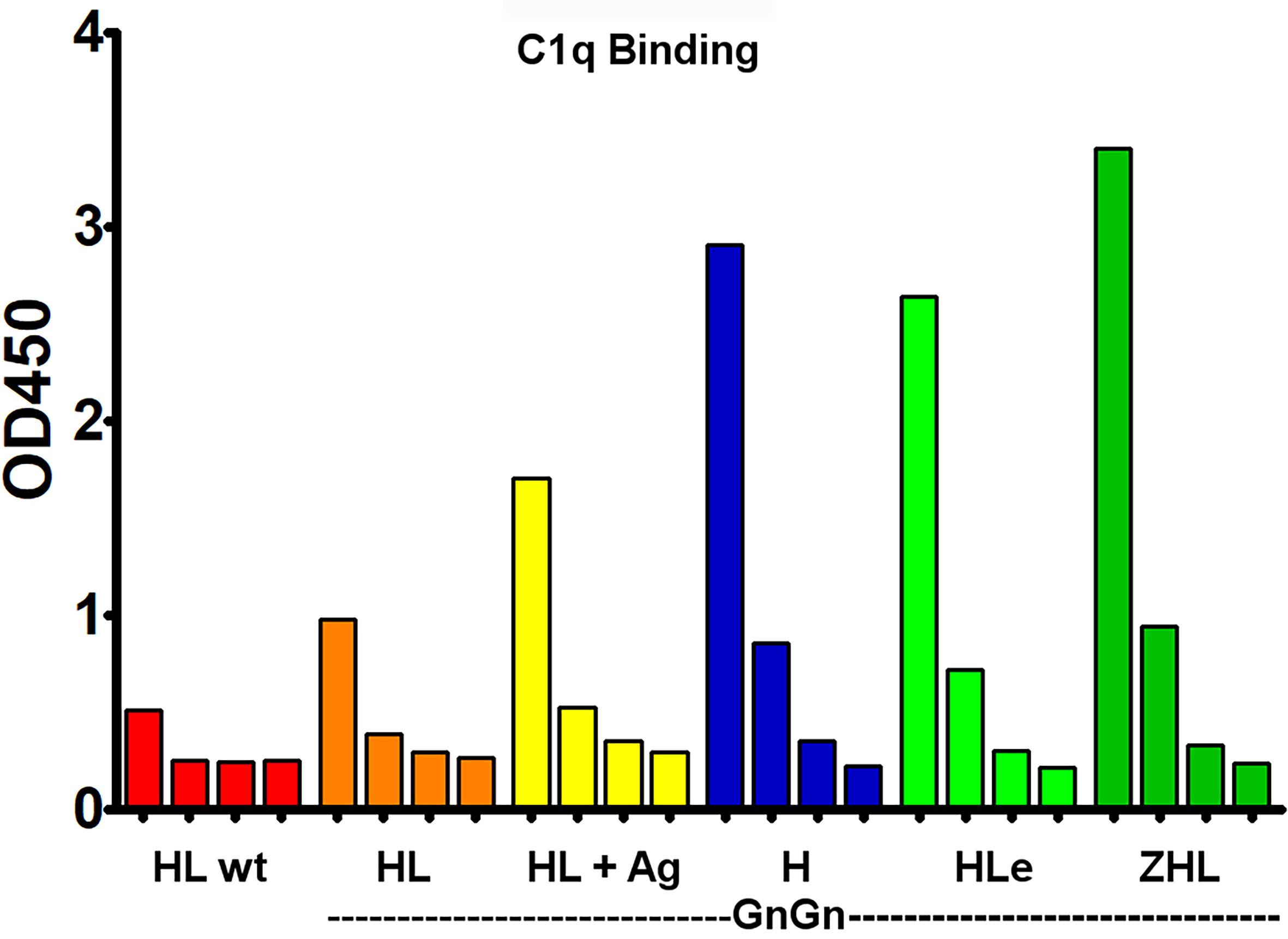

### Supplemental Figure 3

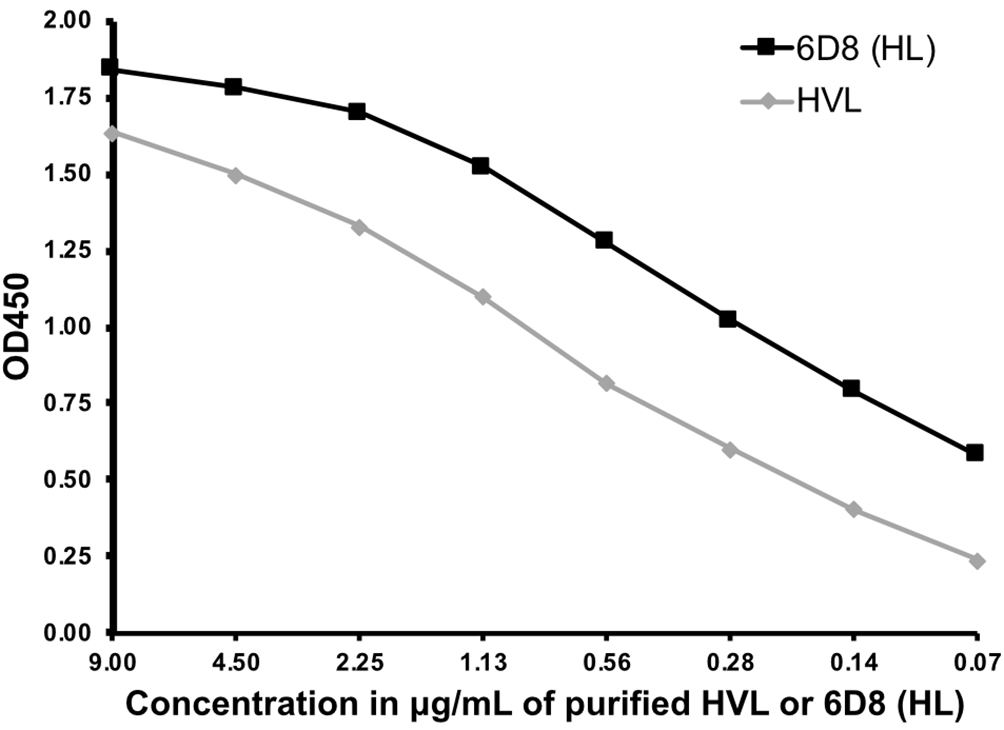

### Supplemental Figure 4

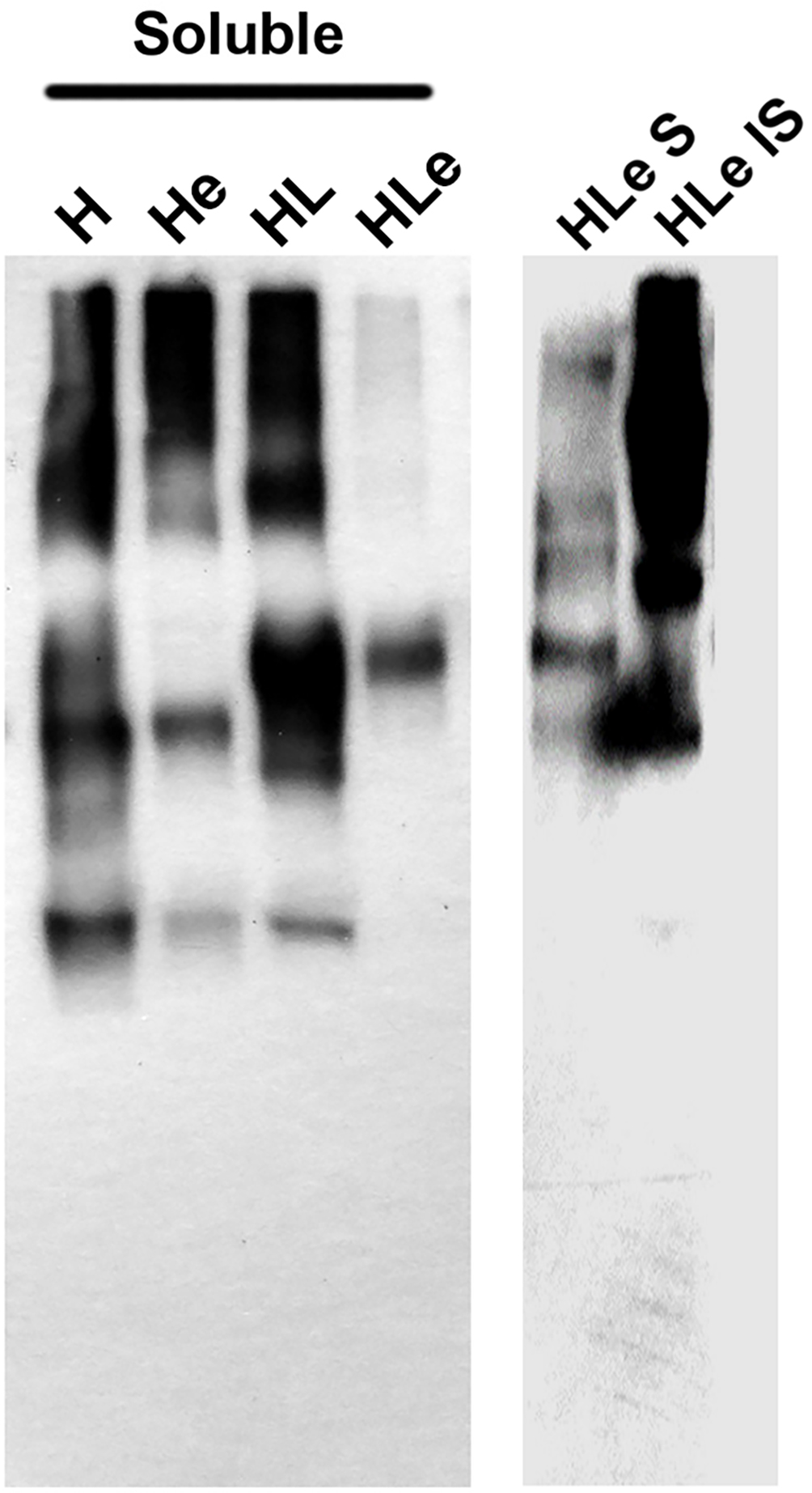

### Supplemental Figure 5

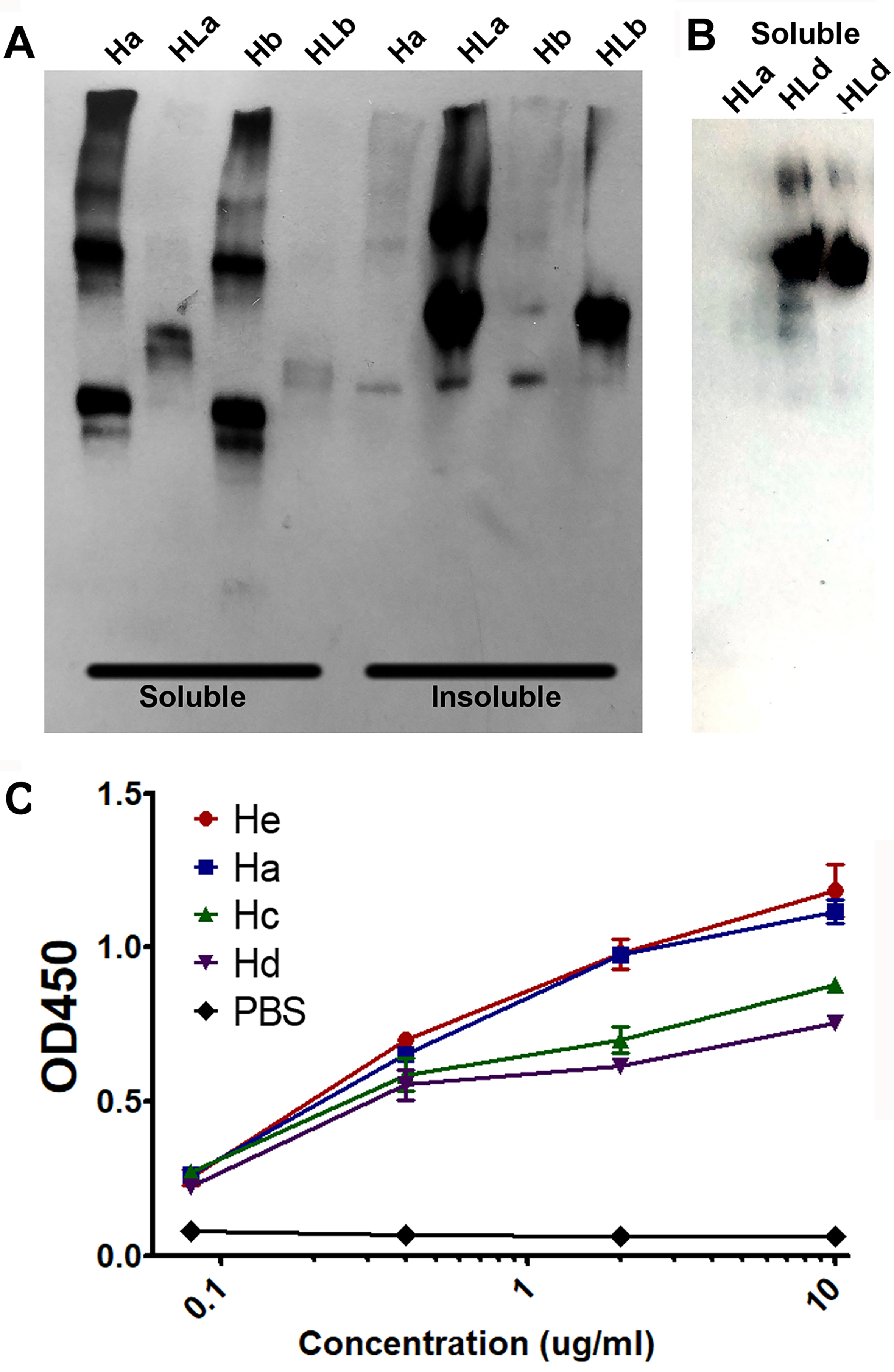
