## Supplemental Methods for "A highly expressing, soluble, and stable plant-made IgG fusion vaccine strategy enhances antigen immunogenicity in mice without adjuvant"

### Supplemental Figures

#### Figure S1. C1q binding comparison between wildtype and glycoengineered plants

ELISA plates were coated with 10 μg/ml human C1q and incubated with 5 μg/ml each purified construct. Constructs were detected using polyclonal goat anti-human IgG-HRP. Mean OD_450_ values from three replicates are shown ± standard error with two stars indicating p < 0.01 as measured by one-way ANOVA with comparisons between the indicated groups. Abbreviations: wt, constructs made in wildtype *N. benthamiana* plants; GnGn, constructs made in glycoengineered plants silenced for xylosyltransferase and fucosyltransferase.

#### Figure S2. C1q binding comparison between IgG fusions

ELISA plates were coated with 10 μg/ml human C1q and incubated with 10-fold serial dilutions of each purified construct starting at 5 μg/ml. Constructs were detected using polyclonal goat anti-human IgG-HRP. Mean OD_450_ values from three replicates are shown. Abbreviations: wt, constructs made in wildtype *N. benthamiana* plants; GnGn, constructs made in glycoengineered plants silenced for xylosyltransferase and fucosyltransferase; Ag, dengue cE tagged with the 6D8 epitope; ZHL, same as HL but with ZE3 fused to the N-terminus of the 6D8 heavy chain.

#### Figure S3. Epitope binding of single chain 6D8

ELISA plates coated with 900 ng of purified epitope-tagged protein were incubated with serial dilutions of either HVL or full-length 6D8 antibody. The bound constructs were detected with HRP-labeled mouse anti-human IgG (Fc-only) antibody and the absorbance was read at OD_450._ Abbreviations: HVL, single-chain antibody with the variable heavy chain linked to the variable light chain region that is fused to the 6D8 heavy chain constant region; 6D8, full-length antibody containing both heavy and light chains.

#### Figure S4. RIC insolubility

Protein was extracted from leaves of *N. benthamiana* agroinfiltrated with the indicated constructs and separated by SDS-PAGE followed by western blotting under nonreducing conditions. The constructs were detected using goat anti-human IgG-HRP. Soluble (S) refers to the clarified crude leaf extract, while insoluble (IS) refers to the pellet following clarification that had been resuspended in SDS sample buffer.

#### Figure S5. Solubility and binding of 6D8 epitope tag mutants

(A, B) Protein was extracted from leaves of *N. benthamiana* agroinfiltrated with the indicated constructs and separated under nonreducing conditions by SDS-PAGE followed by western blotting using goat anti-human IgG-HRP as probe. Soluble refers to the clarified crude leaf extract, while insoluble refers to the pellet following clarification that had been resuspended in SDS sample buffer. The epitope mutant designated “a” contains epitope sequence VYKLDISEA; the epitope mutant designated “b” contains epitope sequence VYKLDISE; the epitope mutant designated “c” contains epitope sequence YKLDISE; the epitope mutant designated “d” contains epitope sequence YKLDIS. (C) ELISA plates coated with serial dilutions of purified heavy chains containing the indicated epitope tag mutants were probed with full-size 6D8 followed by goat anti-human kappa-HRP. Mean OD_450_ values are shown from three replicates ± standard error.
